# Differential expression of cyclins mRNA in neural tissues of BoHV-1- and BoHV-5- infected cattle

**DOI:** 10.1101/674002

**Authors:** Maia Marin, Mercedes Burucúa, Daniel Rensetti, Juan José Rosales, Anselmo Odeón, Sandra Pérez

## Abstract

**Introduction:** Bovine alphaherpesvirus types 1 (BoHV-1) and 5 (BoHV-5) are closely related alphaherpesviruses. BoHV-5 causes non-suppurative meningoencephalitis in calves. BoHV-1 is associated with several syndromes and, occasionally, can cause encephalitis. Although both viruses are neurotropic and they share similar biological properties, it is unknown why these alphaherpesviruses differ in their ability to cause neurological disease.

**Materials and Methods:** Neural tissue samples were collected from BoHV-1- and BoHV-5-intranasally inoculated calves during acute infection, latency and reactivation. The levels of cyclins mRNA expression in neural tissue from calves infected with BoHV-1 or BoHV-5 were analyzed by qRT-PCR. Data were analyzed by Relative Expression Software Tool (REST).

**Results:** Striking differences in the levels of cyclins mRNA were observed between uninfected and infected tissues, particularly in trigeminal ganglion (TG). During acute infection, higher levels of cyclin A2, E1 and B1 were observed in BoHV-1 and BoHV-5-infected TG compared with uninfected TG. mRNA levels of cyclins A2 and E1 were downregulated in olfactory cortex. During latent infection with BoHV-1 and BoHV-5, cyclin A2 and E1 were downregulated in olfactory cortex and cervical medulla whereas cyclin B1 was upregulated in BoHV-1-infected olfactory and frontal cortex and in cervical medulla after BoHV-5 infection. A marked increase of cyclins A2 and E1mRNA levels was detected in TG of BoHV-5-latently-infected cattle. Unlike in uninfected TG, in BoHV-1 and BoHV-5-infected TG, cyclin B1expression was detectable. During reactivation, the levels of cyclin A2, B1 and E1 mRNA increased in TG. The expression levels of cyclins in TG during BoHV-5 latency suggest that these viruses utilize different strategies to persist in the host.

**Conclusion:** Bovine alphaherpesviruses neuropathogenicity might be influenced by the differential control of cell cycle components by these herpesviruses. This is the first report on BoHV-5 modulation of cyclins expression in neural tissues from its natural host.

## Introduction

Bovine alphaherpesvirus types 1 (BoHV-1) and 5 (BoHV-5) are closely related alphaherpesviruses. BoHV-5 is the causal agent of non-suppurative meningoencephalitis in calves [1], a condition which is highly prevalent in South America, particularly Argentina and Brazil. BoHV-1 is associated with several syndromes in cattle, including respiratory disease, abortions, genital disorders [2] and, occasionally, encephalitis [3,4]. Although both viruses are neurotropic and they share similar biological properties, it is unknown why these alphaherpesviruses differ in their ability to cause neurological disease.

The infectious cycle of alphaherpesviruses is characterized by stages of acute infection, latency and reactivation. Sensory neurons within the trigeminal ganglion (TG) are the main site of latency. Latent virus can sporadically reactivate under conditions of natural stress or by glucocorticoid administration. These episodes of reactivation lead to virus excretion and dissemination to susceptible hosts [5]. While BoHV-1 reactivation is subclinical, reactivation of BoHV-5 may occur in the presence of mild neurological signs [1,6].

Many viruses interact with host factors that regulate cell cycle progression in order to facilitate their own replication [7]. The cell cycle can be divided in four phases: G1, S (DNA synthesis), G2 and M (mitosis). Regulation of these phases relies on the formation of complexes between cyclins and cyclin-dependent kinases (cdk) which catalyze the progression of the cell cycle [8] by activating or inactivating target proteins. There are quality control points (checkpoints) that ensure the fidelity of cell division, by controlling that each step of the cycle is completed before the next one takes place [9]. Cells enter into S phase influenced by cyclin E and later by cyclin A2. Maximal activity of both cyclins occurs during this phase. Cyclin A activity continues into G2 and cyclin B1accumulation controls the entry and exit of the cell from M phase [10]. In terminally differentiated neurons, the cell cycle must be kept in constant control since re-initiation of the cycle leads to neuronal death [11]. Studies on BoHV-1 infection have shed light on some strategies of the virus to subvert the control of the cell cycle for its own purpose [12,13]. On the other hand, there is no information on BoHV-5 interactions with cell cycle components. The latency-related (LR) gene of BoHV-1is abundantly expressed in latently-infected neurons and it is known to participate in the modulation of the cell cycle [12,14]. The LR gene of BoHV-1 and BoHV-5 are substantially different [15]. Thus, it is possible that they differ in the modulation of key regulatory molecules involved in cell cycle progression and this might be important for BoHV neuropathogenesis. Thus, the levels of cyclins mRNA expression in neural tissue from calves infected with BoHV-1or BoHV-5 were analyzed. This is the first report on BoHV-5 modulation of cyclins expression in neural tissues from its natural host.

## Materials and methods

In this study, neural tissue samples were collected from BoHV-1- and BoHV-5-intranasally inoculated calves. Fourteen BoHV-free and seronegative cross-bred, 1 year-old calves were used. In Group 1 (primary acute infection; *n* = 4), two calves were inoculated with 10^6.3^ TCID_50_ of BoHV-1 and the other two with a similar inoculum of BoHV-5. They were euthanased at 6 days post-infection (dpi). Group 2 (latency; *n* = 4) consisted of two calves inoculated with (10^3^ TCID_50_) of BoHV-1 or BoHV-5 and they were euthanased at 24 dpi. In Group 3 (reactivation; *n* = 4), two calves were inoculated with 10^3^ TCID_50_ of BoHV-1 and two with BoHV-5. At 20 dpi they received an intravenous dose of 0.1 mg/kg dexamethasone (Dexametona, Schering Plough) and two intramuscular doses 24 and 48 h later. They were euthanased at 25 dpi. Group 4 (mock-infected; *n* = 2) calves were inoculated with culture medium. One calf was euthanased at 6 dpi and the other received dexamethasone under the same regime as calves in Group 3. This calf was euthanased at 25 dpi. All procedures for animal handling and experimentation were performed according to the Animal Welfare Committee of the University of the Center of Buenos Aires Province (Res. 087/02). At 6 dpi calves in BoHV-1 and BoHV-5-infected group were excreting virus at high titers in nasal and ocular secretions, demonstrating that this was the peak of the acute infection period. Furthermore, viral DNA was detected in the neural tissue samples collected at this time-point. At 24 dpi virus was not isolated from secretions or TG. However, viral DNA was present in TG samples from infected calves at this time point. Thus, establishment of latency was confirmed. The results of these experimental inoculations have been previously published [16–20].

Total RNA from tissue samples was extracted using TRIzol reagent (Invitrogen, Carlsbad, CA, USA) according to the manufacturer’s protocol and digested with DNase I Amplification Grade (Invitrogen, Carlsbad, CA, USA). Complementary DNA (cDNA) was synthesized and negative controls, omitting the RNA or the reverse transcriptase, were included. Real-time RT-PCR reactions for cyclins A2, B1 and E1 were performed using 800 nM specific primers (Table 1), 1X PCR Master Mix (FastStart Universal SYBR Green Master Rox, Roche, Mannhein, Germany) and 1 μl of cDNA in a final volume of 20 μl.

**Table 1.**
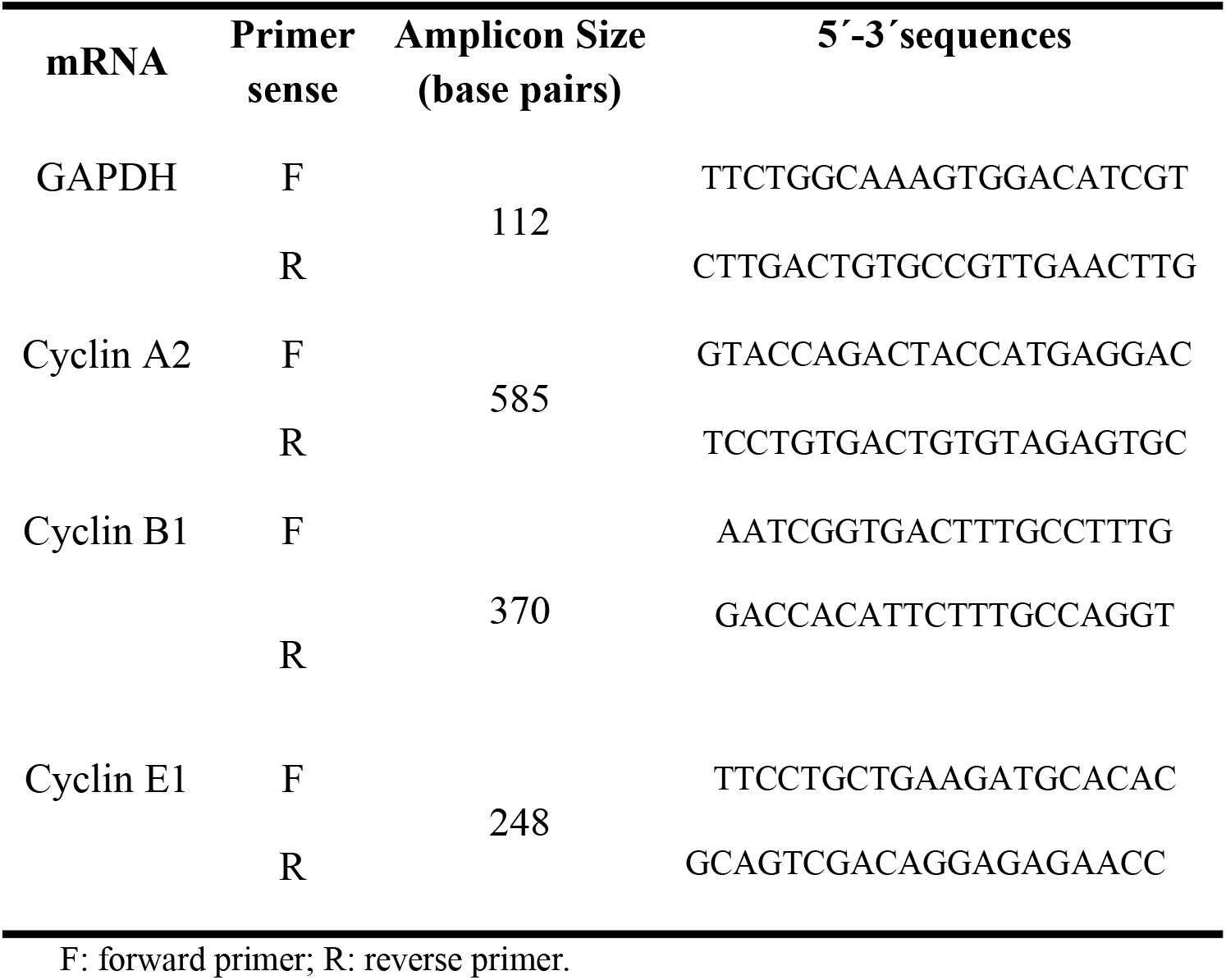
Sequences of primers for mRNA relative quantification by Real time RT-PCR

Amplifications were carried out in an Applied Biosystems 7500 cycler. Expression of the housekeeping gene glyceraldehyde-3-phosphate dehydrogenase (GAPDH) was used as control [21]. Samples were run in duplicate and negative controls for cDNA synthesis and PCR procedures were included. After amplification, a melting curve analysis was performed, which resulted in a single product-specific melting curve. The amplification efficiency was determined for each gene using 10-fold dilutions of the cDNA. The results are reported as the mean fold change of cyclin transcription levels in tissues from infected animals over the levels detected in mock-infected calves. The relative expression analysis was performed using the Relative Expression Software Tool (REST, Qiagen Inc., Valencia, CA, USA).

## Results

Striking differences in the levels of cyclins mRNA expression were observed between uninfected and infected tissues (Table 2).

**Table 2.** Relative expression of bovine cyclins A2, E1and B in nervous system of BoHV-1- and BoHV-5-infected calves

| Gene | Tissue | Acute infection |  |  | Latency |  |  | Reactivation |  |  |
| --- | --- | --- | --- | --- | --- | --- | --- | --- | --- | --- |
|  |  | Control | BoHV-1 | BoHV-5 | Control | BoHV-1 | BoHV-5 | Control | BoHV-1 | BoHV-5 |
| CYCLIN A2 | Olfactory cortex | 1 | 0.01* | 0.02 | 1 | 0.00* | 0.00* | 1 | 0.88 | 1.62 |
|  | Frontal cortex | 1 | 0.59 | 2.00 | 1 | 2.19 | 2.66 | 1 | 0.46 | 0.59 |
|  | Posterior cortex | 1 | 0.90 | 1.38* | 1 | 2.80* | 8.30 | 1 | 0.08* | 0.03* |
|  | Cervical medulla | 1 | 6.68 | 0.26* | 1 | 0.20 | 0.07 | 1 | 0.06* | 0.23 |
| CYCLIN E1 | Olfactory cortex | 1 | 0.00* | 0.02* | 1 | 0.00* | 0.00* | 1 | 0.84 | 1.98 |
|  | Frontal cortex | 1 | 0.58 | 1.53 | 1 | 1.97* | 3.98 | 1 | 2.13 | 3.98* |
|  | Posterior cortex | 1 | 1.08 | 1.63 | 1 | 3.80* | 26.05 | 1 | 1.08 | 0.71 |
|  | Cervical medulla | 1 | 3.22 | 0.13* | 1 | 0.03* | 0.02* | 1 | 0.11* | 0.37 |
| CYCLIN B1 | Olfactory cortex | 1 | 0.97 | 1.22 | 1 | 5.58* | 3.89 | 1 | 1.25 | 0.64 |
|  | Frontal cortex | 1 | 1.19 | 2.68 | 1 | 4.52* | 4.55 | - | + | + |
|  | Posterior cortex | 1 | 1.08 | 1.03 | 1 | 3.40 | 3.98 | - | + | + |
|  | Cervical medulla | 1 | 7.32 | 0.69* | 1 | 4.40 | 11.8* | 1 | 0.11* | 2.35* |

The results represent the mean fold change of gene transcription levels in the areas of the nervous system of infected calves over levels detected in tissue sections of uninfected calves, which served as the control group. +: Gene cDNA was detected in the nervous tissue sample analyzed; -: Gene cDNA was not detected in the nervous tissue sample analyzed; \*: statistically significant differences ( $P < 0.05$ ) with respect to the uninfected control for this tissue sample.

These differences were particularly evident in TG at all stages of the infectious cycle. Cyclin B1 was undetected in TG of uninfected calves. During acute infection, cyclin A2 and E1, were significantly higher (*p* < 0.05) in BoHV-1 and BoHV-5-infected TG compared with uninfected TG. Cyclin B1 was undetected in acutely-infected TG. mRNA levels of cyclins A2 and E1 were downregulated in olfactory cortex (*p* < 0.05) (Table 2). At this stage of the infectious cycle, cyclin E1 and B were downregulated in cervical medulla of BoHV-5-infected calves (*p* < 0.05) (Table2). Interestingly, in BoHV-1-infected TG, cyclin A2 and E1 mRNA levels were 23.7 and 2.6 fold-higher than the levels detected in BoHV-5 infected TG (Fig 1).

**Fig 1.**
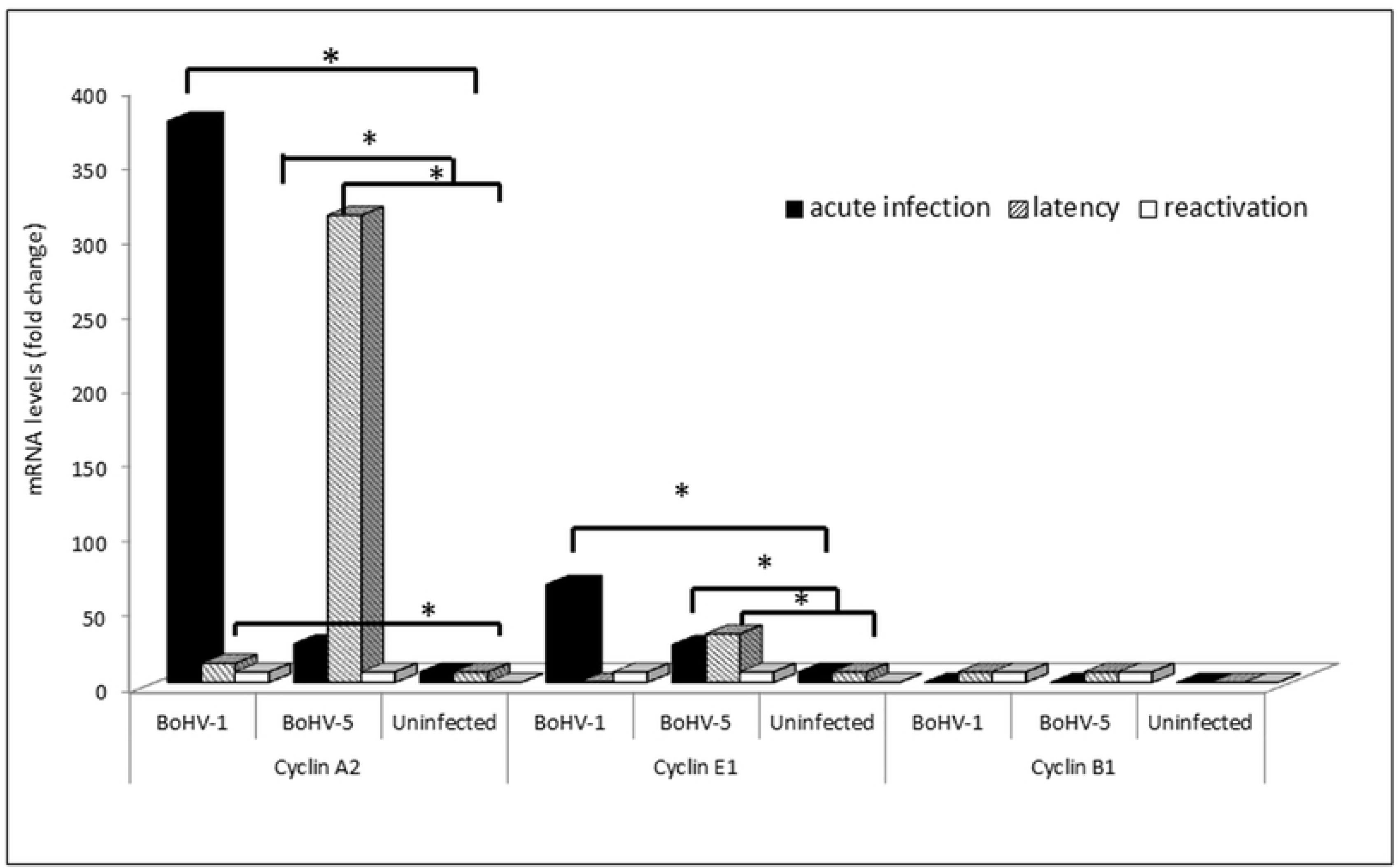
Cyclins A2, B1 and E1 mRNA expression in BoHV-infected trigeminal ganglion. The results represent the mean fold change of cyclins A2, B1 and E1 expression levels in trigeminal ganglion of BoHV-1 and BoHV-5-infected calves over levels detected in the same tissue of uninfected calves, which served as the control group. *: statistically significant differences (*P* <0.05).

However, significant differences were not detected with respect to BoHV-5-infected TG (*p* > 0.05). During latent infection with BoHV-1 and BoHV-5, a significant downregulation (*p* < 0.05) of cyclin A2 and E1 was observed in olfactory cortex and cervical medulla and only a slight but significant increase (*p* < 0.05) was detected in frontal (cyclin A2 and B1) and posterior cortex (cyclins A2 and E1). Cyclin B1 expression was significantly higher (*p* < 0.05) in cervical medulla of BoHV-5 infected calves (Table 2). On the contrary, at this stage, a marked increase in cyclins A2 and E1 mRNA levels was detected in TG of BoHV-5-infected cattle. Cyclin A2 and cyclin E1 levels were 114 and 85.7 fold-higher, respectively, in BoHV-5- compared with BoHV-1-infected TG (*P*<0.05). Cyclin B1 mRNA expression was detected during latent infection of BoHV-1 or BoHV-5 (Fig 1). Even during reactivation, although unquantifiable, cyclins A2, B and E1 mRNA levels were upregulated when compared with levels in uninfected TG (Fig 1). Furthermore, during reactivation, a significant decrease (*p* < 0.05) in cyclin A2 mRNA expression was detected in cervical medulla of BoHV-1- and posterior cortex of BoHV-1 and BoHV-5-infected animals. A slight increase (*p* < 0.05) in cyclin E1 and cyclin B levels were recorded in frontal cortex and cervical medulla, respectively, of BoHV-5-infected calves (Table 2).

## Discussion

In this study, remarkable differences in the modulation of cyclins expression by BoHV-1 and BoHV-5 were detected. This was particularly striking in TG, the main site of latency of alphaherpesviruses. Peak cyclins A2 and E1 expression was evident in TG of BoHV-1-acutely-infected calves. Since in highly differentiated cells, such as neurons, cyclin A expression induces apoptosis [22], these findings correlate with the strategies of the virus to enhance productive infection [13]. On the contrary, in BoHV-5-infected TG, peak mRNA of both cyclins was detected during latency. It is known that BoHV-1 LR gene products interact with cyclin A [14]. Inhibiting the expression of cell cycle components and the consequent neuronal apoptosis during latency is essential for virus survival within the host [12,13]. Similar to the findings of our study, in a rabbit model, Schang et al. [12] demonstrated that BoHV-1 induced cyclin A expression in TG during acute infection and reactivation. In a previous work, we found that, compared to BoHV-1, BoHV-5 induces lower levels of neuronal apoptosis in TG and this correlated with lower levels of viral replication [18]. It is also likely that these levels of BoHV-5 replication correlate with the lower levels of cyclin A2 at this stage. Both viruses induced the expression of cyclin B1 in TG during latency and reactivation. Normally, entry into mitosis is tightly regulated and requires the cyclin B1/cdk1 complex to reach threshold levels of activity [23]. In our study, cyclin B1 was upregulated in TG during latency of both viruses, in the olfactory and frontal cortex of BoHV-1-infected calves and in cervical medulla of BoHV-5-infected cattle. Upregulation of cyclin B1 expression correlates with neurons being driven into G2/M transition. These abnormal alterations in differentiated cells end into abortive events, leading to degenerating neurons and neuropathology [24]. Although for BoHV-5 infection significant differences in the levels of cyclin B1 expression with respect to uninfected tissues were not observed, it is also likely that these levels have biological significance in BoHV-5 neuropathogenesis.

Overall, the high expression levels of cyclins in TG during BoHV-5 latency suggest that these viruses utilize different strategies to persist in the host and this might be related to differential functional activities of BoHV-5 LR gene. Further studies, including the analysis of cyclin protein levels, are necessary to understand the precise mechanism of alphaherpesviruses regulation of the cell cycle. Nevertheless, these findings suggest that neuropathogenecity might be influenced by the distinct control of cell cycle components.

## Acknowledgements

We especially thank María Rosa Leunda and Susana Pereyra for their contribution and support to the study.

## Author contributions

**Conceptualization:** Sandra Pérez

**Data curation:** Maia Marin, Mercedes Burucúa, Juan José Rosales, Daniel Rensetti

**Formal analysis:** Sandra Pérez, Maia Marin, Anselmo Odeón

**Investigation:** Sandra Pérez, Maia Marin, Anselmo Odeón

**Methodology:** Maia Marin, Mercedes Burucúa, Juan José Rosales, Daniel Rensetti

**Project administration:** Sandra Pérez

**Supervision:** Sandra Pérez, Maia Marin

**Validation:** Sandra Pérez, Maia Marin

**Writing-original draft:** Sandra Pérez

**Writing-review and editing:** Sandra Pérez, Maia Marin

